# Laminar Dynamics of Target Selection in the Posterior Parietal Cortex of the Common Marmoset

**DOI:** 10.1101/2023.08.15.553425

**Authors:** Janahan Selvanayagam, Kevin D. Johnston, Stefan Everling

## Abstract

The lateral intraparietal area (LIP) plays a crucial role in target selection and attention in primates, but the laminar microcircuitry of this region is largely unknown. To address this, we used ultra-high density laminar electrophysiology with Neuropixels probes to record neural activity in the posterior parietal cortex (PPC) of two adult marmosets while they performed a simple visual target selection task. Our results reveal neural correlates of visual target selection in the marmoset, similar to those observed in macaques and humans, with distinct timing and profiles of activity across cell types and cortical layers. Notably, a greater proportion of neurons exhibited stimulus related activity in superficial layers whereas a greater proportion of infragranular neurons exhibited significant post-saccadic activity. Stimulus-related activity was first observed in granular layer putative interneurons, whereas target discrimination activity emerged first in supragranular layers putative pyramidal neurons, supporting a canonical laminar circuit underlying visual target selection in marmoset PPC. These findings provide novel insights into the neural basis of visual attention and target selection in primates.

## Introduction

At any given moment, we are faced with many more stimuli than can be processed simultaneously. To cope with this limitation, the process of attention acts to filter irrelevant stimuli and preferentially select those relevant for the guidance of behaviour. In foveate animals such as primates, visual attention and eye movements are closely linked, and the neural mechanisms underlying these processes and their relation to one another has been a topic of intensive investigation. Convergent evidence from anatomical, lesion, fMRI, TMS, and neurophysiological studies has demonstrated that attention and eye movements are supported by an extensively interconnected and largely overlapping network that includes the frontal eye fields (FEF) within prefrontal cortex, the lateral intraparietal area (LIP) within the posterior parietal cortex (PPC), and the midbrain superior colliculus (SC), an area critical for the generation of eye movements (see for review Johnston & Everling, 2008; McDowell et al., 2008).

The role of LIP in attentional and oculomotor control has been a topic of considerable interest, owing in part to its anatomical interposition between sensory and motor areas. LIP receives extensive inputs from multiple visual cortical areas, and as noted above is reciprocally interconnected with FEF and SC (Andersen et al., 1990; Baizer et al., 1991; Lewis & Van Essen, 2000; Lynch et al., 1985; Schall, 1995). As such, it has been conceptualized as a transitional link between visual processing and saccade generation. Consistent with this, single neurons in LIP have been shown to exhibit both visual and saccade related responses (Andersen et al., 1987).

More direct evidence has been provided by studies in macaque monkeys trained to perform variants of the visual search task, in which a target stimulus is selected from an array of distractors. Pharmacological inactivation of LIP has been shown to induce deficits in visual search performance (Wardak et al., 2002). Neurophysiological studies have revealed that the activity of LIP neurons evolves to discriminate targets from distractors presented within their response fields in advance of saccades to the target location (Ipata et al., 2006; Mirpour et al., 2009; Thomas & Paré, 2007), and that the time of this discrimination is predictive of the reaction times of targeting saccades (Thomas & Paré, 2007). Thus, the activity of LIP neurons may be said to instantiate a process of saccade target selection, in which an initial stage of visual selection is followed by activity related to the forthcoming saccade.

Broadly speaking, for tasks requiring target selection, the activity of LIP neurons resembles closely that of areas to which it projects. Neurons both in FEF (Thompson et al., 1996) and SC (McPeek & Keller, 2002; Shen et al., 2011) discriminate targets from distractor stimuli and discharge in advance of saccades. Although activity in both of these areas (Dorris et al., 1997; Hanes & Schall, 1996; Paré & Hanes, 2003) has been more directly linked to saccade initiation than that in LIP (Gottlieb & Goldberg, 1999), the considerable overlap in discharge properties across areas invites detailed investigations of the intrinsic mechanisms shaping the selection process within each area which in turn regulate the signals sent between them to fully understand their respective contributions to target selection. Anatomical and physiological evidence has demonstrated that area LIP possesses separate output channels to the FEF and SC. Cortico-cortical projections exhibit a visual bias and originate predominately in layers II/III, while corticofugal projections originate exclusively in layer V and exhibit a bias toward saccade-related activity (Ferraina et al., 2002; Lynch et al., 1985; Schall, 1995). To date, the laminar dynamics shaping these activity differences remain poorly understood, and although canonical circuit models have provided theoretical accounts with respect to visual cortex (Douglas & Martin, 1991) and the FEF (Heinzle et al., 2007) few studies have investigated directly the laminar flow of information by conducting simultaneous recordings across cortical layers (but see Bastos et al., 2018; Godlove et al., 2014; Nandy et al., 2017; Ninomiya et al., 2015; Pettine et al., 2019). The flow of neural activity in the primate posterior parietal cortex Is unknown.

The lack of laminar recordings in fronto-parietal networks is due in large part to the practical difficulty in accessing areas such as FEF and LIP in macaques due to their locations deep within sulci. In contrast, the common marmoset monkey (*Callithrix jacchus*) has a relatively lissencephalic cortex, making it well-suited for such investigations. Recent work has identified homologous regions to macaque and human FEF and LIP in marmosets using a variety of methods, including cyto- and myeloarchitectural features, anatomical connections, resting state functional connectivity, task-based fMRI activations, intracortical microstimulation, and single-unit electrophysiology (Collins et al., 2005; Feizpour et al., 2021; Ghahremani et al., 2017, 2019; Ma et al., 2020; Reser et al., 2013; Rosa et al., 2009; Schaeffer et al., 2019; Selvanayagam et al., 2019). Here, we addressed the knowledge gap in the understanding of laminar dynamics and their role in instantiating the process of saccadic target selection by carrying out laminar electrophysiological recordings in the posterior parietal cortex of marmosets using ultra-high density Neuropixels probes (Jun et al., 2017) while they performed a simple visual target selection task in which they generated saccades to a target stimulus presented in either the presence or absence of a distractor. We observed neural correlates of visual target selection similar to those observed in macaques and humans, the timing of which varied across neuron type and cortical layer.

## Results

### Behavioural Performance

Marmosets performed visually guided saccades in a simple target selection task wherein blocks of “single-target” and “distractor” trials were presented to the animal (see Figure 1a).

**Fig 1.**
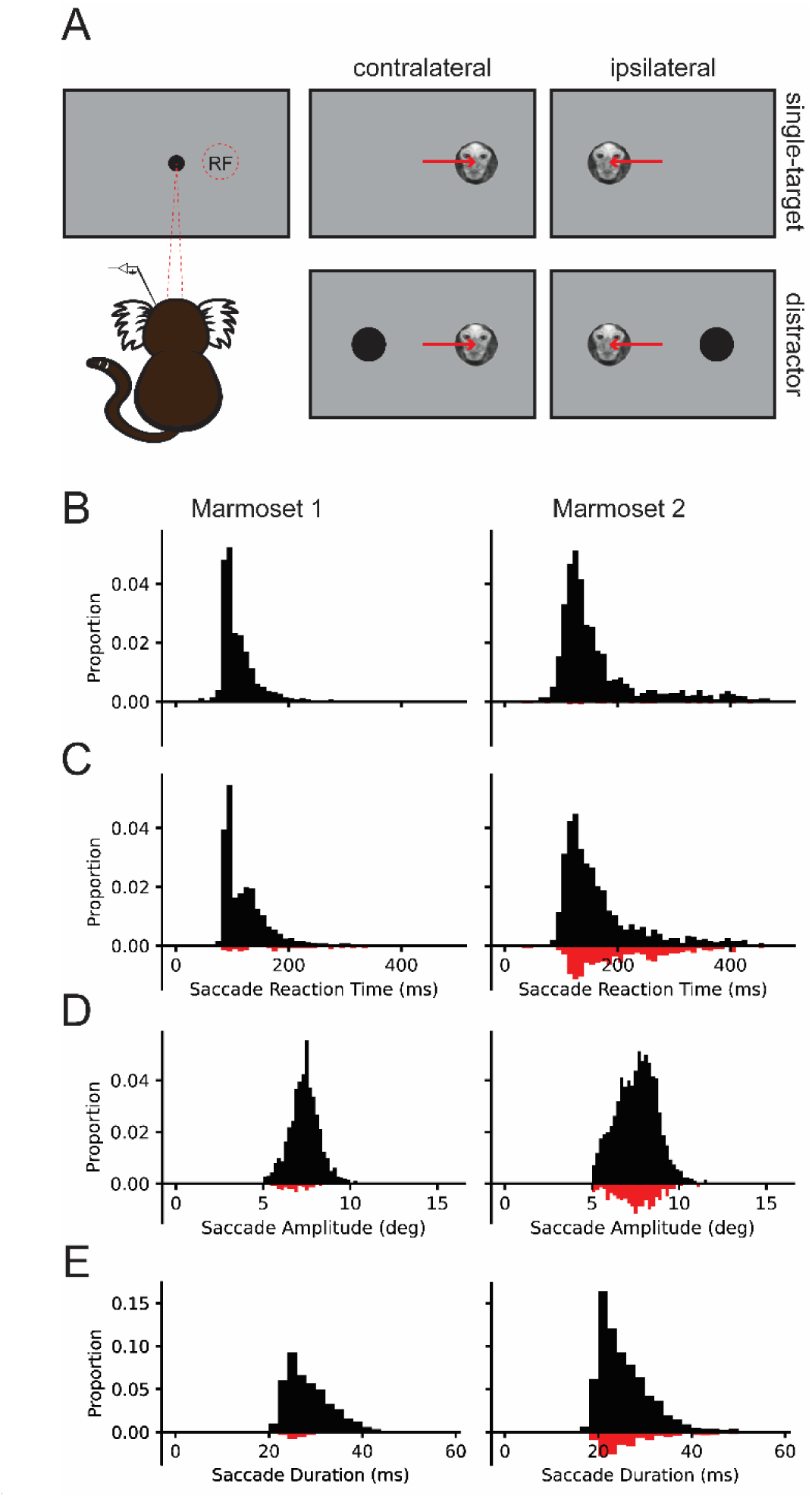
Task design and behavioural performance. (A) Schematic representing task design for “single-target” and “distractor” trials, where the target falls in (contralateral) or out of the receptive field (RF). (B) Saccade reaction time histograms for “single-target” and “distractor” trials for each animal separately. Saccade amplitude (C) and duration (D) histograms for each animal separately across all conditions.

Animals were required to fixate on a central fixation stimulus (0.5° radius black circle on a grey background) for 300-500 ms at the beginning of each trial. On “single-target” trials, a single target (1° diameter image of a marmoset face) was presented 6° to the left or right of the fixation stimulus and subjects were required to make a saccade to the target to obtain a viscous liquid reward of acacia gum. On “distractor” trials, a distractor stimulus (0.5° radius black circle) was simultaneously presented in the opposite hemifield. Trials in which no saccade at least 4° in amplitude was made were marked as “no response” and were not included in further analysis.

Trials in which saccades were made to the target were marked as correct and trials in which saccades landed anywhere else were marked as incorrect. We conducted 22 recording sessions, 8 in Marmoset M and 14 in Marmoset N, in which animals performed 162-438 trials (*M*=248.7 trials). Accuracy was significantly lower on “distractor” trials (Mean ± SEM; Marmoset M: 89.9 ± 2.2%; Marmoset N: 74.0 ± 4.0%, see Figure 1b) than on “single-target” trials (Marmoset M: 100.0 ± 0.0%, Marmoset N: 96.4 ± 0.5%, see Figure 1c), Marmoset M: *t*(7)=4.57, *p*=.003, Marmoset N: *t*(13)=5.82, *p*<.001, and median saccade reaction times (SRT) were significantly longer, Marmoset M: *t*(7)=3.29, *p*=0.013, Marmoset N: *t*(13) = 3.79, *p*=.002, (Marmoset M: “distractor”=110.0±4.0ms vs “single-target”=99.4±1.6ms; Marmoset N: “distractor”=146.8±6.5ms vs “single-target”=139.0±5.2ms). Saccade amplitude and durations did not differ significantly between conditions nor on correct vs incorrect trials (all *p*’s > .05; see Figure 1d-e). Taken together, these results reveal a distractor- induced reduction in performance suggesting an additional stage of processing on these trials.

### Determining recording locations, cortical layers, and putative neuron classes

To determine recording locations we acquired high resolution, anatomical T2 images from each animal. Prior to scanning, a custom-designed grid with 1.5mm diameter holes spaced at 1mm was inserted in the animals’ recording chambers and filled with iodine solution. The filled grid holes provided landmarks for determining the locations of identified areas within the recording chamber. We then aligned these images to a high-resolution ex-vivo MRI template (REF?) aligned with a group RS-fMRI functional connectivity map of the SC (https://www.marmosetbrainconnectome.org, Schaeffer et al., 2022). We identified a region of strong functional connectivity in the PPC corresponding to the location of area LIP (see Figure 2a-b; Ghahremani et al., 2019; Schaeffer et al., 2019). Marmosets subsequently underwent aseptic surgeries in which we opened trephinations of approximately 3 mm in diameter over this region.

**Fig 2.**
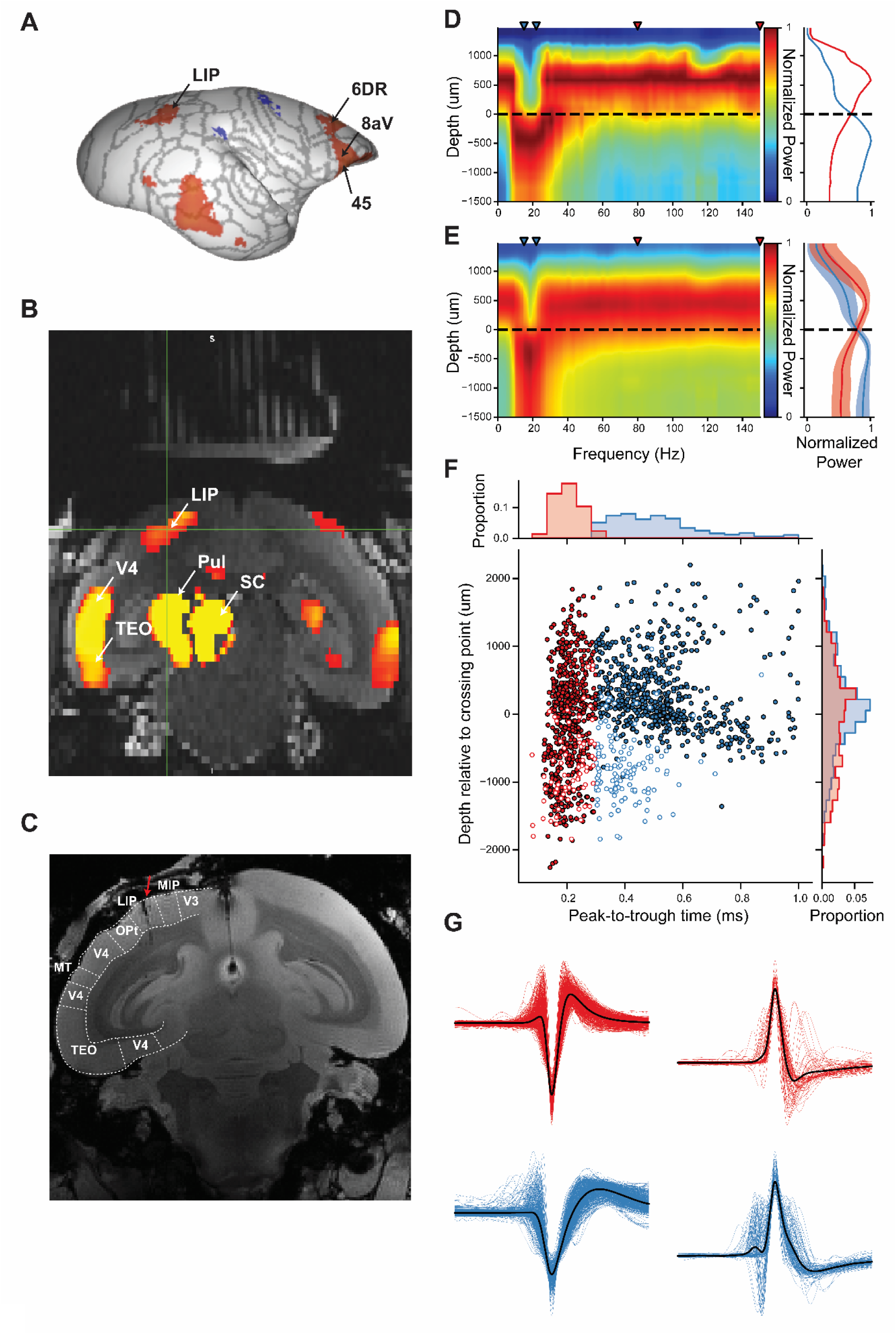
Localization of recording locations, layer assignment and cell type classification. (A) Surface map of resting state functional magnetic resonance imaging (RS-fMRI) functional connectivity (FC) with superior colliculus to identify lateral intraparietal area (LIP). (B) Coronal slice of anatomical MRI of Marmoset M with an overlay of FC maps from A interpolated to native space of Marmoset M to identify location of LIP relative to the grid. (C) Ex-vivo anatomical MRI of Marmoset N with Paxinos et al., (2012) boundaries overlaid confirming electrode tract locations (as indicated by red arrow) in LIP. LFP power aligned to stimulus onset across depths and frequencies (left) and normalized power in selected ranges (right; blue: 15-22 Hz, red: 80-150 Hz) are shown for an example session (D) and the average of all sessions (E). The crossing point between lower and higher frequencies is marked by a dotted line. LIP = Lateral intraparietal area, MIP = Medial intraparietal area, TEO = temporal area TE occipital part, MT = middle temporal area, OPt = occipito-parietal transitional area

We conducted 26 electrode penetrations in two animals (Marmoset M: 8 penetrations in 8 sessions; Marmoset N: 18 penetrations in 14 sessions) in which we advanced either one or two Neuropixels electrodes (Jun et al., 2017) in this region and recorded the activity of 1366 well-isolated single neurons. For each penetration, we determined cortical layers by identifying the crossover point between the power spectral density (PSD) of low (15-22 Hz) and high (80-150 Hz) frequency ranges in the local field potentials (LFP) across depths as done in previous work (Mendoza-Halliday et al., 2023) (see Figure 2c). Based on visual inspection of the distribution of isolated neurons distributed along the length of the electrode shank, and the known density of neuronal distributions within the cortical layers in this region of marmoset cortex, we classified all neurons that fell within 200 µm below to 300 µm above as being in granular Layer IV, and all others as supragranular or infragranular. To classify putative interneurons and pyramidal cells, the established approach of using the peak-trough duration was employed (Ardid et al., 2015; Hussar & Pasternak, 2012; McCormick et al., 1985; Mitchell et al., 2007) (see Figure 2d).

Interestingly, a large proportion of neurons with positive spiking waveforms were observed (198, 14.5 %), which were largely restricted to the broad waveforms observed in deeper layers. For these waveforms, we inverted the waveform before evaluating the peak-trough duration.

### Evaluating stimulus and saccade-related responses in LIP neurons

To identify task-modulated neurons, we computed the mean discharge rates from 50 ms after stimulus onset to 25 ms after saccade onset for conditions and compared it to the mean baseline activity 200 ms before stimulus onset. Examining the conditions separately, 319 (23.35%) neurons were significantly modulated in the “single target” contralateral condition as compared to 112 (8.2%) in the “single-target” ipsilateral condition; for the “distractor” trials, 329 (24.08%) were significantly modulated when the target was presented in the contralateral hemifield as compared to 262 (19.18%) when the distractor was presented in the contralateral hemifield. Overall, pooling across conditions, a total of 390 (28.55%) neurons exhibited significant modulations in discharge rates during task performance (see Figure 3). The proportion of modulated neurons per layer and putative cell class were as follows: supragranular (BS: 26.98%, NS: 35.20%), granular (BS: 29.14%, NS: 35.03%), and infragranular (BS: 23.15%, NS: 25.09%). For these neurons, we conducted Pearson R correlations to determine whether this activity correlated with the SRTs for contralateral and ipsilateral trials separately; the discharge activity of 32 (8.2%) and 33 (8.5%) neurons were significantly correlated with SRTs (*p*’s < .05) respectively, suggesting little correspondence between the activity of these neurons and SRTs.

**Fig 3.**
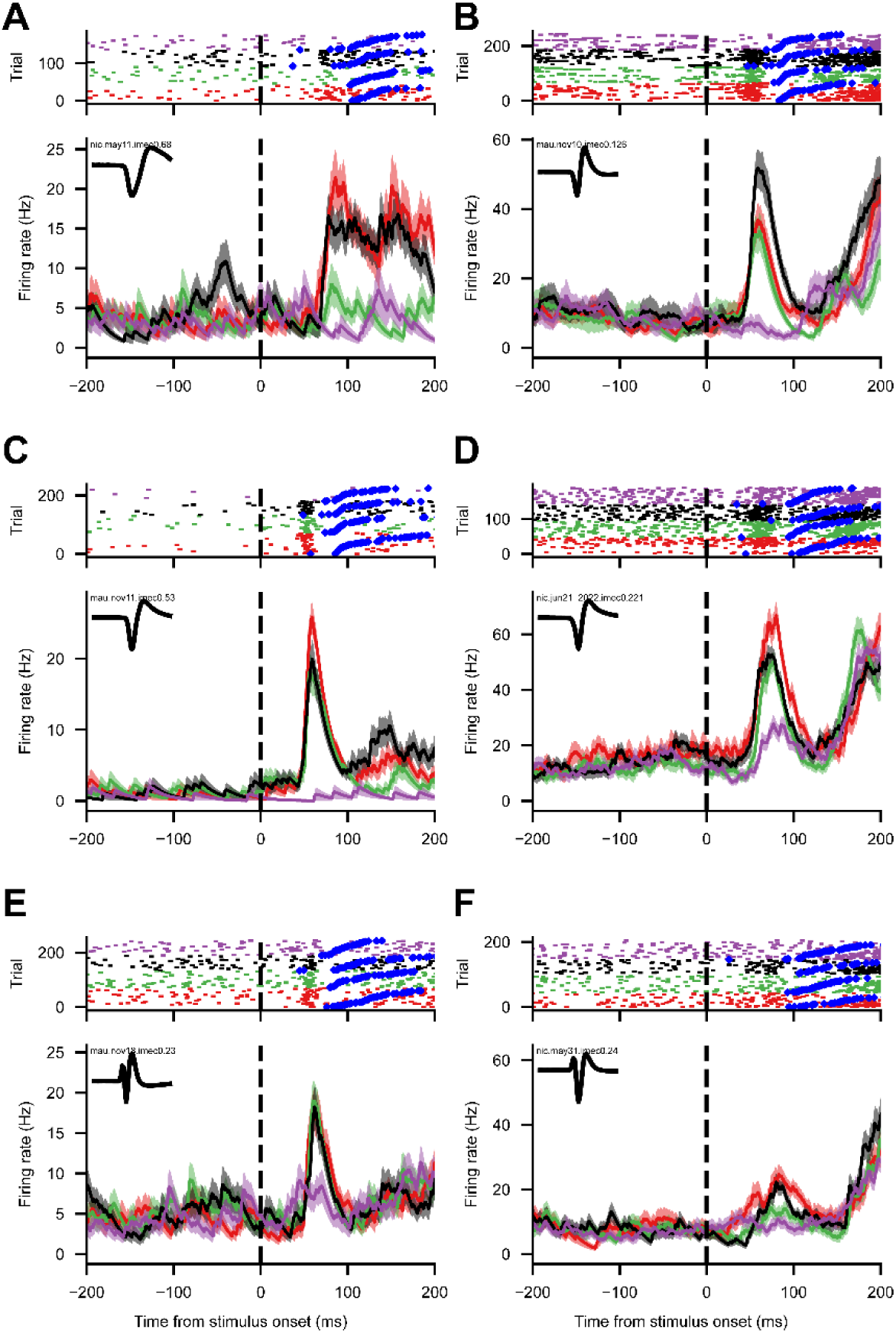
Example visual neurons. Raster plots and spike density functions (SDF) aligned to stimulus onset for example broad (A, C, E) and narrow (B, D, F) -spiking neurons from supragranular (A, B), granular (C, D) and infragranular (E, F) layers with visual activity. Black = target contralateral, Red = target contralateral & distractor ipsilateral, Purple = target ipsilateral, Green = target ipsilateral & distractor contralateral. Blue diamonds represent saccade onset. Mean waveform in inset SDF figure.

We then compared the activity on distractor trials in which the target was in the contralateral or ipsilateral hemifield. One-hundred and sixty-eight (12.3%) neurons significantly discriminated between targets and distractor presented in the contralateral hemifield, of which 135 (80% of discriminating neurons) showed greater activity for the target stimulus (see Figure 4); supragranular (BS: 10.71%, NS: 14.80%), granular (BS: 16.55%, NS: 15.29%), infragranular (BS: 5.91%, NS: 10.75%). For these neurons, to assess the magnitude of target discrimination, we conducted Receiver Operating Characteristics (ROC) analyses (Green & Swets, 1966) comparing the distributions of activity on the trials in which the preferred (i.e., the stimulus with the greater mean discharge activity in the task epoch) or non-preferred stimulus was presented in the contralateral hemifield. We computed auROC values on discharge rates within successive 15 ms intervals from stimulus onset to 200 ms after stimulus onset. We then determined the magnitude and time from stimulus onset of the maximal auROC value for each neuron. Medians across layers and putative cell class were as follows: supragranular (BS: .602, 76 ms; NS: .598, 78 ms), granular (BS: .583, 94 ms; NS: .645, 83 ms), infragranular (BS: .613, 100 ms; NS: .599, 77ms). Notably, the maximal auROC values were generally observed before the median SRTs, however, the timing and magnitude of the discrimination did not differ appreciably between layers and cell types.

**Fig 4.**
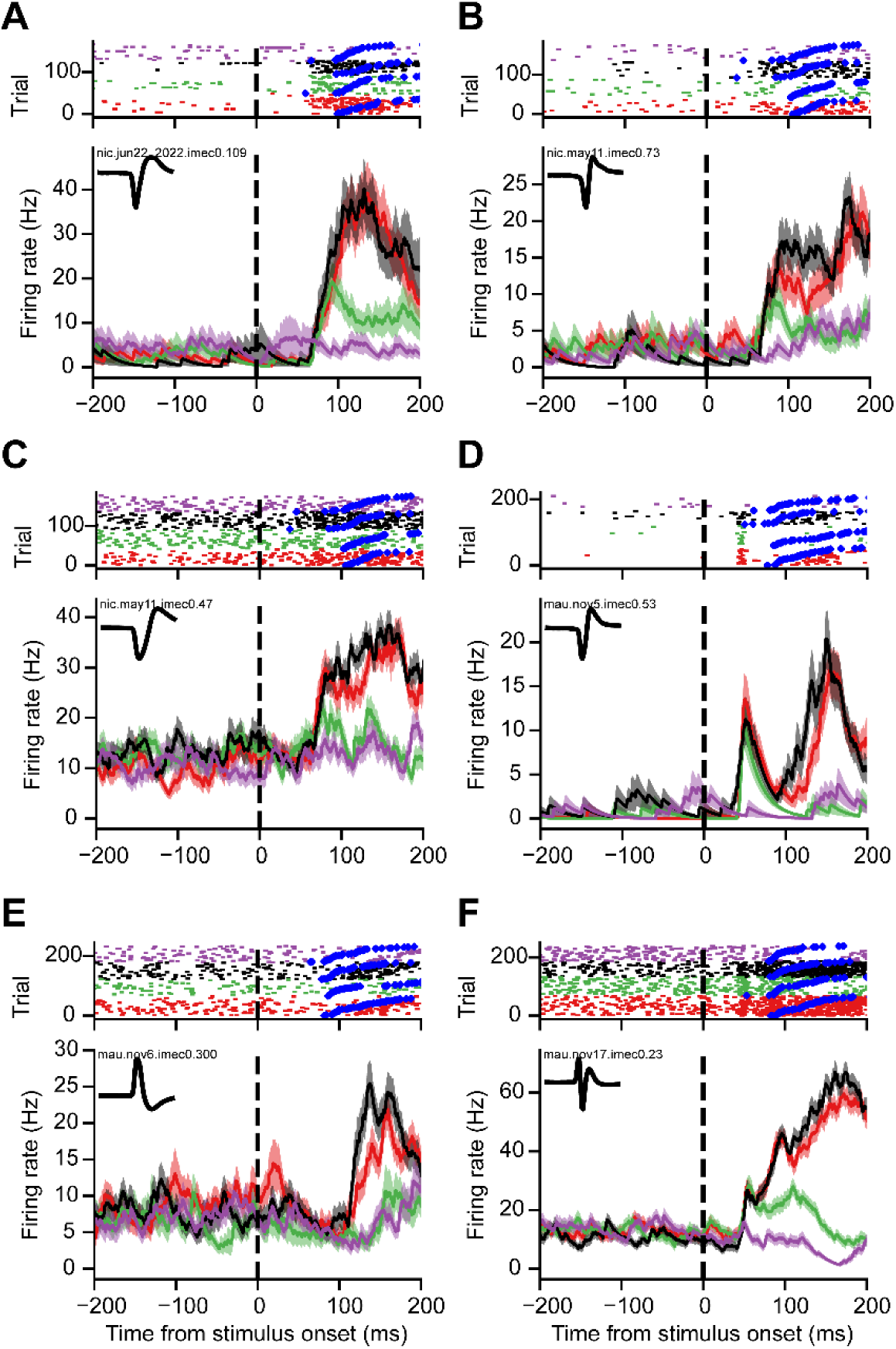
Example target discriminating neurons. Raster plots and spike density functions (SDF) aligned to stimulus onset for example broad (A, C, E) and narrow (B, D, F) -spiking neurons from supragranular (A, B), granular (C, D) and infragranular (E, F) layers with activity discriminating between target and distractor stimuli. Black = target contralateral, Red = target contralateral & distractor ipsilateral, Purple = target ipsilateral, Green = target ipsilateral & distractor contralateral. Blue diamonds represent saccade onset. Mean waveform in inset SDF figure.

We additionally observed a large proportion of neurons that displayed strong post-saccadic modulations in activity across conditions. Generally, this activity started at saccade offset, peaked approximately 50-100 ms later and often persisted for 300-500 ms. To identify neurons with significant post-saccadic activity, we computed the mean discharge rates from 50- 150ms after saccade offset where we observed the peak of the activity and compared it to the 200 ms prestimulus baseline used above, separately for each condition. For correct trials, 969 neurons (70.94%) had significant post-saccadic activity in at least one condition, 688 neurons in at least two conditions, 391 in at least three conditions, 203 in all four conditions; 551-581 neurons for each condition; see Figure 5. Post-saccadic activity did not appear to correspond with stimulus-related activity; of the 329 neurons with significant stimulus-related activity in the contralateral “single-target” condition 58 neurons had significant post-saccadic activity in the ipsilateral “single-target” condition, 71 in the contralateral and 101 in both. For the “distractor” conditions, we examined post-saccadic activity on error trials and observed that only half the number of neurons had significant post-saccadic activity (ipsilateral: 231 neurons as compared to 551, 119 neurons in both; contralateral: 288 neurons as compared to 566, 165 in both). In sum, a large proportion of neurons exhibited post-saccadic activity and this activity varied depending on stimulus identity and task performance.

**Fig 5.**
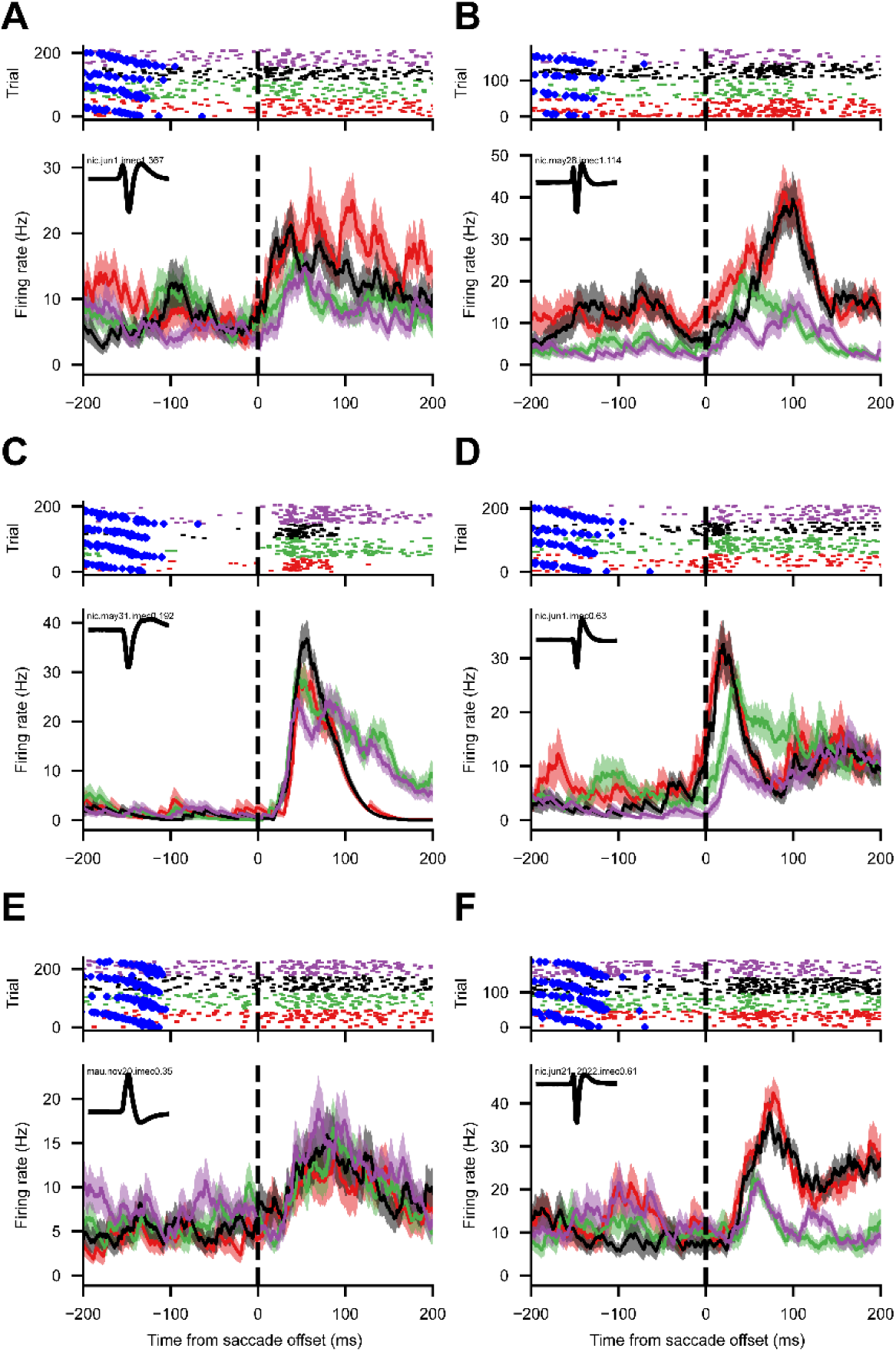
Example post-saccadic neurons. Raster plots and spike density functions (SDF) aligned to saccade offset for example broad (A, C, E) and narrow (B, D, F) -spiking neurons from supragranular (A, B), granular (C, D) and infragranular (E, F) layers with significant post- saccadic activity. Black = target contralateral, Red = target contralateral & distractor ipsilateral, Purple = target ipsilateral, Green = target ipsilateral & distractor contralateral. Blue diamonds represent stimulus onset. Mean waveform in inset SDF figure.

For comparison with the above, we determined the proportion of neurons with significant post-saccadic activity across conditions. We then conducted a logistic regression to investigate the effects of layer, cell-type, and epoch (task, discrimination, post-saccadic) on the likelihood that a neuron has significantly different discharge activity. This model explained significantly more variance than the reduced two-way models (*p* < .05), and revealed that NS infragranular neurons were less likely to be significantly modulated in the task interval, and BS infragranular neurons were less likely to significantly discriminate between target and distractor but were more likely to have significant post-saccadic activity as compared to respective granular and supragranular layer neurons (*p*’s < .05).

In sum, we observed in marmosets LIP neurons which were significantly modulated in a visual target selection task and, in particular, those that discriminated between target and distractor stimuli before making a saccade. Further, this activity was observed across cortical laminae and cell types, albeit in slightly different proportions; supragranular and granular neurons were more likely to demonstrate stimulus and target selection related activity whereas infragranular neurons were more likely to have significant post-saccadic activity.

### Stimulus related activity first emerges in narrow spiking granular layer neurons

To examine if and how the emergence of stimulus-related activity differs across cortical layers and cell types, we investigated the population activity using generalized additive models (GAMs). GAMs are a type of statistical model which fits data to a “smooth” curve comprised of many line segments by estimating the value at each “knot”, the boundaries of these segments (Hastie & Tibshirani, 1986). In this manner, GAMs can capture complex, non-linear relationships such as how neural activity varies over time (Cadarso-Suarez et al., 2006). Here, for the entire population of recorded neurons, we modelled the odds of a spike at each point in time as a function of time from stimulus onset, the cortical layer (supragranular, granular and infragranular) and cell type (BS, NS) for the contralateral and ipsilateral “single-target” condition. This model significantly improved fit as compared to the reduced models (*p* < .05).

As the vast majority of neurons only exhibited significant increases in discharge activity for contralateral as compared to ipsilateral “single-target” trials, we could evaluate the onset of stimulus-related activity by comparing the activity between these conditions. We computed difference smooths between contralateral and ipsilateral conditions with a 99.9% CI, identified time points where this difference smooth deviated from 0 (see Figure 6), and determined the earliest time point where stimulus-related activity was observed for each cell type and cortical layer. Stimulus-related activity first emerged in NS granular and BS supragranular neurons (35ms), followed by NS supragranular neurons (37 ms) and finally in BS granular layer neurons (42 ms). In sum, this suggests that stimulus-related activity first emerges in the granular layer, then in supragranular layers and occurs first in NS, i.e., putative interneurons. The population stimulus-related activity in infragranular layers did not reach significance at any time point.

**Fig 6.**
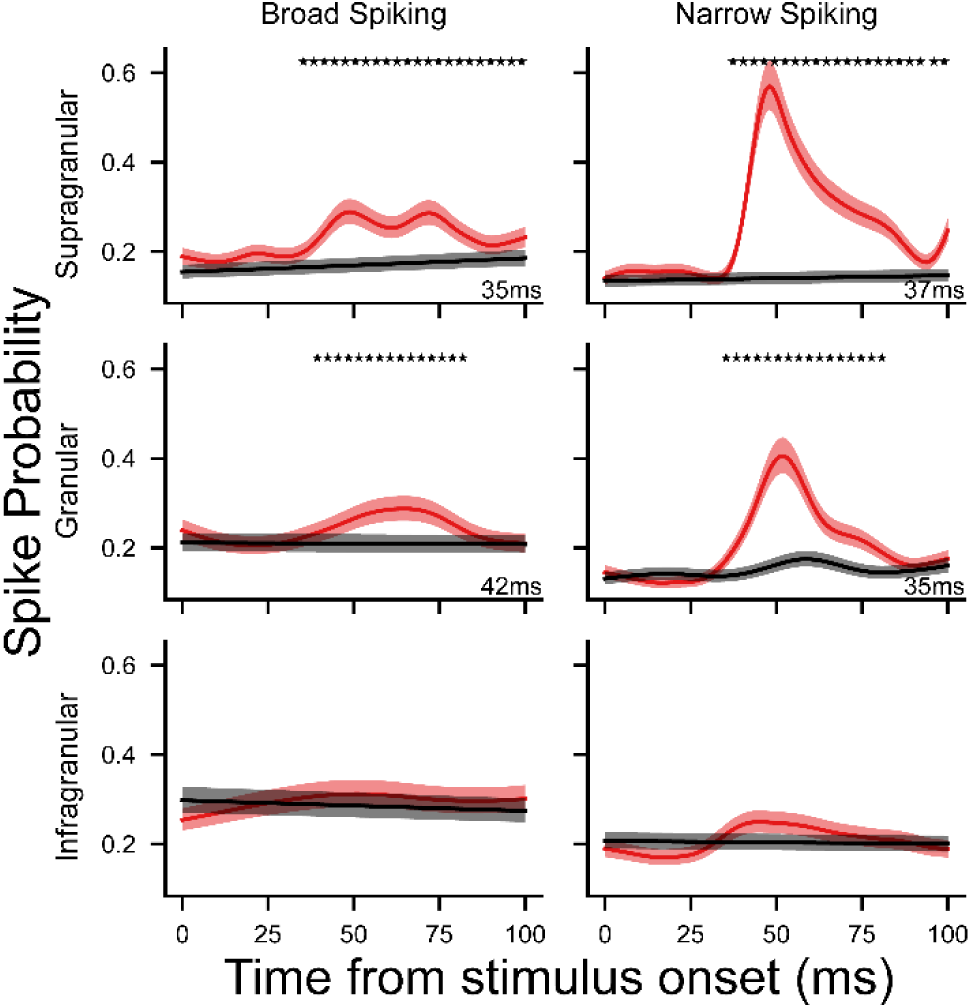
Generalized additive model fit for population stimulus-related activity. Odds of a spike at a given point in time are estimated using the time from stimulus onset, the putative layer the neuron is found in (supragranular, granular, or infragranular), putative cell class (NS or BS), and the condition of the given trial (“single target” ipsilateral or contralateral) with trial and neuron as random effects. Spike probability in the ipsilateral (black) and contralateral (red) conditions are plotted here for broad (left) and narrow (right) spiking neurons for supragranular, granular and infragranular layers. * = significant difference between conditions at 99.9% CI. First significant time point noted in bottom right corner in ms.

### Target discrimination related activity first emerges in broad spiking supragranular neurons

Next, we examined how target discrimination activity first emerges in the population activity using a GAM where we modelled odds of spiking using time, cortical layer, cell type and condition (ipsilateral vs contralateral “distractor” trials; *p* < .05). We then computed difference smooths between the conditions with a 99.9% CI, identified time points where this difference smooth deviated from 0 (see Figure 7), and determined the earliest time point where target discrimination activity was observed for each cell type and cortical layer. We observed target discrimination activity first in BS supragranular neurons (35 ms), followed by infragranular neurons (NS: 36 ms, BS: 37 ms), then in BS granular layer neurons (39 ms), then in NS supragranular neurons (41 ms) and finally in NS granular layer neurons (44ms). Altogether, we see target discrimination emerges rapidly in superficial layers and predominantly in BS neurons.

**Fig 7.**
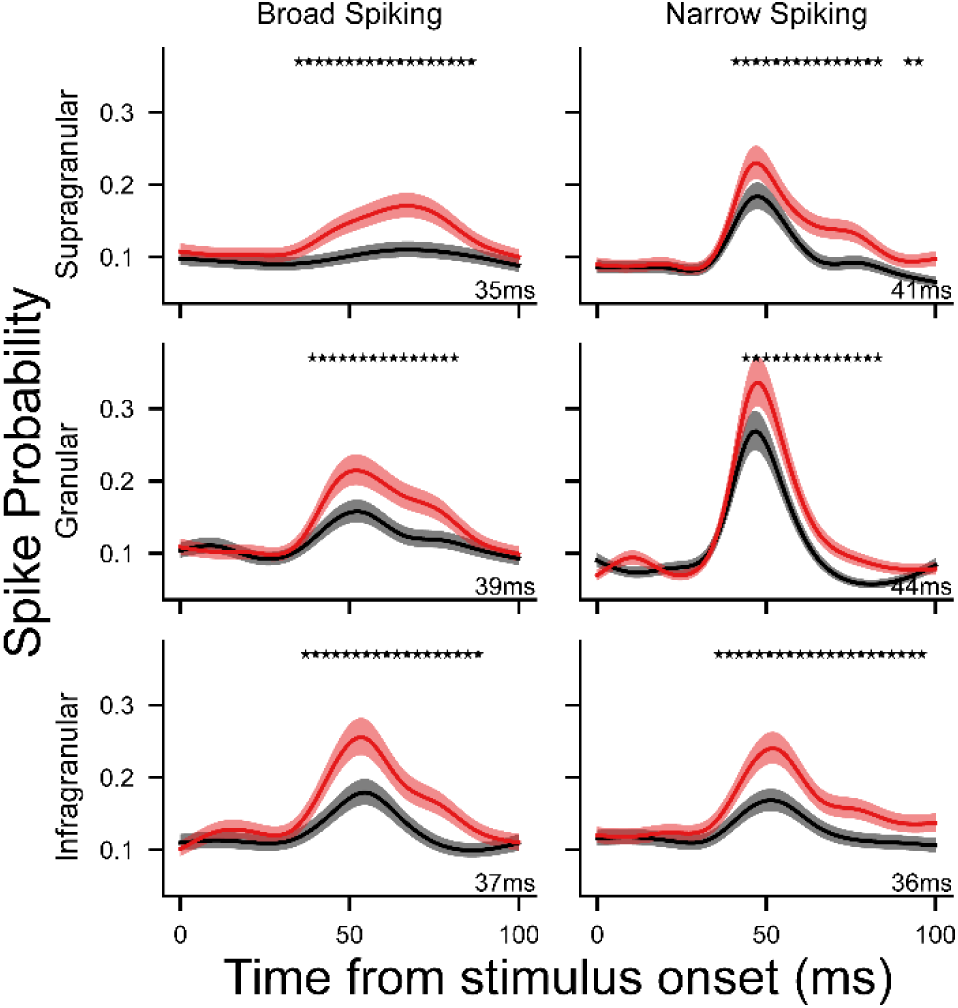
Generalized additive model fit for population target discrimination activity. Odds of a spike at a given point in time are estimated using the time from stimulus onset, the putative layer the neuron is found in (supragranular, granular, or infragranular), putative cell class (NS or BS), and the condition of the given trial (“distractor” preferred vs non-preferred) with trial and neuron as random effects. Spike probability in the preferred (red) and non-preferred (black) conditions are plotted here for broad (left) and narrow (right) spiking neurons for supragranular, granular and infragranular layers. * = significant difference between conditions at 99.9% CI. First significant time point noted in bottom right corner in ms.

In sum, although neurons with stimulus-related and target discrimination activity were observed across cortical laminae, subtle differences in the timing of this activity were observed, suggesting the granular layer as the primary input and supragranular layers as the first to discriminate between targets and distractors.

## Discussion

The laminar microcircuitry underlying visual target selection and saccade control in PPC remains poorly understood due to limitations of previously used animal models and experimental approaches. Here, we employed ultra-high density laminar electrophysiology in the PPC of common marmosets as they completed a saccadic target selection task to address this gap. As expected, we observed neurons with stimulus-related activity and, for the first time in the marmoset, neurons that discriminated between target and distractor stimuli. The stimulus-related activity observed here first emerged in the granular layer, followed by supragranular layers, with population activity in infragranular layers never reaching significance. This activity emerged first in putative interneurons followed by putative pyramidal neurons. Conversely, activity discriminating between target and distractor stimuli first emerged in supragranular neurons, followed by infragranular and finally granular layers, usually first appearing in putative pyramidal neurons. Altogether, the observed patterns support the existence of a canonical circuit consistent with previous models (Douglas & Martin, 2004; Heinzle et al., 2007).

Since its first description in (Andersen et al., 1987; Barash et al., 1991a, 1991b), LIP has been the focus of intensive investigation for its role in the control of visual attention and eye movements. Single neuron recordings in macaque LIP have demonstrated that neurons in this area respond selectively to relevant visual stimuli and are critical in guiding visual attention and saccadic eye movements (Andersen et al., 1987; Barash et al., 1991a, 1991b; Colby et al., 1996; Gnadt & Andersen, 1988; Kusunoki et al., 2000). Subsequent work, typically employing variants of the visual search task, has demonstrated activity in LIP which evolves to discriminate the presence of targets or distractors within their response fields (Ipata et al., 2006; Mirpour et al., 2009; Thomas & Paré, 2007). Investigations using pharmacological interventions and cortical cooling have further demonstrated a causal role for LIP in regulating visual salience (Chen et al., 2020; Wardak et al., 2002). Consistent with these observations, for the first time in the common marmoset, we observed a number of neurons that in a simple target selection task, responded to visual stimuli, a large proportion of which discriminated targets from distractors.

Further, this discrimination activity generally peaked in advance of the upcoming saccade to the target location, consistent with a visual selection process preceding saccade generation. The magnitude of this discharge activity however did not correlate with the SRTs, though this is not surprising as the activity of LIP neurons does not strictly predict the motor plan underlying upcoming saccadic eye movements, but rather represents the current locus of attention across the visual field (Bisley & Goldberg, 2003; Goldberg et al., 2002; Kusunoki et al., 2000).

Surprisingly, a large proportion of neurons displayed significant post-saccadic activity across conditions which began immediately after saccade offset and often persisted up to 500 ms. As this activity is observed for both ipsilateral and contralateral trials in the “single-target” condition, it is unlikely to reflect remapping signals for a stimulus passing through the future receptive field of a neuron, as is observed in LIP neurons for “double-step” saccade paradigms (Duhamel et al., 1992). This activity could reflect the efference copy of the saccade i.e. corollary discharge (Sommer & Wurtz, 2008). In FEF, corollary discharge activity can be observed which is relayed from SC by the medial dorsal nucleus of the thalamus (Sommer & Wurtz, 2004, 2006). The observed activity here could be corollary discharge activity from SC in a similar pathway through pulvinar or from FEF. It has been previously observed that this activity in PPC can reflect saccadic error or saccade duration (Munuera & Duhamel, 2020; Zhou et al., 2016, 2018). It is worth noting that for many of these neurons, this activity varied across conditions and as a function of task performance while saccade amplitude and duration did not, suggesting this activity is not merely an efference copy but may encode other task-relevant variables.

While the activity of LIP neurons has not been shown to be tightly linked to saccade initiation, such activity can be observed in other frontoparietal structures such as FEF and SC with which LIP is strongly interconnected. Notably, LIP projections to these areas are largely segregated within distinct cortical laminae; cortico-cortical projections originate primarily in supragranular layers II/III and tend to convey visual information whereas corticotectal projections originate exclusively from infragranular layer V and primarily carry saccade-related information (Ferraina et al., 2002; Lynch et al., 1985; Schall, 1995). Indeed, computational models based on studies of macaque FEF and observed laminar circuits in cat primary visual cortex (Douglas & Martin, 2004; Heinzle et al., 2007) propose layer IV as the input, layers II/III as being responsible for the rule-based allocation of attention, and layer V as the primary output. These observations motivate investigations of laminar dynamics of areas such as FEF and LIP underlying these differences. Although these are challenging to pursue in the macaque due to the location of these areas in sulci prohibiting laminar electrophysiology, the lissencephalic cortex of the marmoset lends itself well to such investigations.

To this end, we used established methods of identifying cortical layers based on the PSD (Mendoza-Halliday et al., 2023) and classifying putative cell classes on the basis of peak-trough widths (Ardid et al., 2015; Hussar & Pasternak, 2012; McCormick et al., 1985; Mitchell et al., 2007). We reliably observed a crossing point in the power of low and high frequencies across depths, indicative of granular layer IV, from which we were able to separate cortex into supragranular, granular, and infragranular layers. Regarding cell type classification, we observed a larger than expected proportion of positive-negative waveforms, which were largely restricted to BS infragranular neurons. These waveforms likely correspond to spikes recorded at the apical dendritic trunk of pyramidal neurons with large apical dendritic arbors which may be more commonly encountered in deeper layers (Boulton et al., 1990, p. 9). For many of these neurons, we observed lower amplitude, negative-positive waveforms on deeper electrode contacts consistent with spikes recorded at the soma. To classify these neurons, we simply inverted the waveform before computing the peak-trough width.

We then assessed how the observed activity varied across cortical laminae and putative cell classes. First, we examined the proportion of neurons with significant stimulus-related, discrimination and post-saccadic activity. NS supragranular/granular layer neurons were more likely to have stimulus-related activity as compared to infragranular neurons. Similarly, superficial BS neurons were more likely to discriminate between targets and distractors.

Conversely, BS infragranular neurons were more likely than their superficial counterparts to display significant post-saccadic activity. These observations are consistent with the proposed role of superficial layers in visual input and attentional deployment and deeper layers for output.

Interestingly, we observed no difference in the maximal magnitude of the discrimination between target and distractor stimuli across layers or putative cell classes. However, the timing of how this activity evolves did differ. We first observed stimulus-related activity in putative interneurons in the granular layer followed by supragranular layer neurons. This is consistent with what is observed in other cortical areas and proposed by theoretical models. Moreover, this is consistent with the anatomy as corticocortical feedforward projections and thalamic input primarily terminate in granular layer IV and to a lesser extent, supragranular layers (Baizer et al., 1991; Matsuzaki et al., 2004). That it is observed first in putative interneurons as compared to pyramidal neurons is perhaps surprising as the primary target of long-range cortical projections are spiny neurons, which are generally pyramidal neurons (Anderson et al., 2011). However, this is characteristic of thalamocortical feedforward inhibition as observed in mouse barrel cortex (Swadlow, 2002). Here it is observed that monosynaptic thalamocortical input to somata of broadly tuned and highly sensitive layer IV interneurons act to rapidly drive inhibition which in turn sharpens the tuning properties of nearby pyramidal cells. Next, also consistent with our hypothesis, we observed discrimination between target and distractor stimuli first in putative pyramidal neurons in supragranular layers. Neurons in this layer are known to share reciprocal projections other key cortical structures involved in visual target selection such as FEF (Ferraina et al., 2002).

In sum, we identified single neurons exhibiting stimulus-related activity and those that discriminate between target and distractor stimuli across all layers and cell types albeit at different proportions and times. These observations are consistent with observations in single neuron investigations of LIP. Ferraina and colleagues (2002) antidromically identified populations of LIP neurons that were either a more superficial cortico-cortical, FEF projecting population, or a deeper corticotectal, SC projecting population. While these populations did possess similar stimulus-related, delay and saccade-related activity, a greater proportion of the more superficial cortico-cortical population exhibited stimulus-related activity whereas a greater proportion of the deeper corticotectal population exhibited delay and saccade-related activity.

These observations are consistent with our own, highlighting a role of more superficial neurons in earlier visual processing and deeper neurons in later saccadic stages. This can also be observed in FEF, where layer V corticotectal neurons represent activity at nearly all stages of visuomotor processing but tended to be more related to movement than more superficial cortico-cortical neurons (Everling & Munoz, 2000; Wurtz et al., 2001). Similarly in V4, a greater proportion of neurons with visual activity and feature selectivity can be observed in superficial layers as compared to a greater representation of eye movement related signals in deeper layers (Pettine et al., 2019; Westerberg et al., 2021).

Altogether, our findings demonstrate single neuron target selection related activity in the posterior parietal cortex of marmoset monkeys. Critically, we found interlaminar dynamics underlying this activity in primate association cortex consistent with a “canonical circuit” resembling that observed in primary visual cortex and proposed for the frontal eye fields. These dynamics are characterized by a flow of neural activity from granular, to supragranular, to infragranular layers, with stimulus-related activity emerging first in granular layer putative interneurons and target discrimination first emerging in supragranular putative pyramidal neurons.

## Methods

### Subjects

Two adult common marmosets (Marmoset M, female, age 22-24 months, weight 328-337 g; Marmoset N, male, age 23-35 months, weight 421-443g) served as subjects in the present study. Prior to these experiments, both animals were acclimated to restraint in two separate custom-designed primate chairs for MRI and electrophysiological experiments which placed them in sphinx and upright positions, respectively. The animals additionally underwent an aseptic surgical procedure in which a combination recording chamber/head restraint was implanted, the purpose of which was to stabilize the head for MRI imaging, eye movement recording, and electrode insertions, and to allow access to cortex for electrophysiological recordings. These procedures have been described in detail previously (Johnston et al., 2018; Schaeffer et al., 2019). All experimental procedures were conducted in accordance with the Canadian Council on Animal Care policy on the care and use of laboratory animals and a protocol approved by the Animal Care Committee of the University of Western Ontario Council on Animal Care. The animals were additionally under the close supervision of university veterinarians throughout all experiments.

### Behavioural training

For training on eye movement tasks, marmosets were seated in a custom primate chair (Johnston et al., 2018) inside a sound attenuating chamber (Crist Instrument Co. Hagerstown MD), with the head restrained. A spout was placed at animals’ mouth to allow delivery of a viscous liquid reward (acacia gum) via an infusion pump (Model NE-510, New Era Pump Systems, Inc., Farmingdale, New York, USA). All visual stimuli were presented on a CRT monitor (ViewSonic Optiquest Q115, 76 Hz non-interlaced, 1600 x 1280 resolution) using either the CORTEX real-time operating system (NIMH, Bethesda, MD, USA) or Monkeylogic (Hwang et al., 2019). Eye positions were digitally recorded at 1 kHz via infrared video tracking of the left pupil (EyeLink 1000, SR Research, Ottawa, ON, Canada).

Marmosets were first trained to fixate on visual stimuli by rewarding 300-600 ms fixations within a circular electronic window with a diameter of 5° centred on circular stimuli consisting of dots with a diameter of 2° presented centrally on the display monitor. Once they were able to perform this subtask reliably, the number of potential fixation locations was increased with the addition of four stimuli presented at +/- 5° abscissa and +/- 5° ordinate. This served both as an initial training stage and allowed us to verify and adjust eye position calibration at the beginning of each experimental session.

Marmosets were then trained on the visual target selection task (see Figure 1a). This task consisted of two trial types. On “single-target” trials, the animals were required to generate a saccade to the location of a single peripheral visual stimulus in order to obtain a liquid reward.

On each trial, they were required to maintain fixation within an electronic window with a diameter of 5° centred on a 0.5° dot presented at the centre of the display monitor for a variable duration of 300-500 ms. Following this, a single target stimulus, a marmoset face (3° diameter), was presented at +/- 6° abscissa. Animals were rewarded for single saccades to the target stimulus which landed within a circular electronic window of 5°, centred on the stimulus. Saccades landing elsewhere were marked as “incorrect”. If no saccade was made within 1 s of target onset, the trial was marked as “no response”. Once marmosets were consistently able to perform 100 or more correct trials of this task within a session, we added an additional “distractor” condition in which a distractor stimulus, a 1° radius black circle, was presented in the opposite hemifield at equal eccentricity to the target stimulus. All fixation and saccade requirements and the timing of trial events was identical to that of single target trials. On distractor trials, single saccades to the target stimulus were rewarded while those made to the distractor location were classified as errors. In the final version of the task the “single target” and “distractor” conditions were run in alternating 20 trial blocks. Marmosets were trained on this task until they could complete 200 trials with at least 70% accuracy in the distractor blocks excluding “no response” trials. At this point we commenced collection of electrophysiological data. The final blocked version of the task including single target and distractor conditions was used for all electrophysiological recording sessions.

### fMRI-Based Localization of Recording Locations

To target LIP for electrophysiological recordings, we conducted an fMRI localizer prior to commencing electrophysiological recordings. To provide landmarks for the location of this area relative to the recording chamber and guide the placement of trephinations allowing access to cortex, a custom-designed in-house printed grid matched to the inside dimensions of the chamber, consisting of 1mm holes at a spacing of 1.5mm, was placed into the chamber and the grid holes filled with iodine solution prior to scanning. This allowed visualization of the chamber and grid coordinates in the MRI images. We then acquired awake anatomical T2 images from each animal and aligned these to a high-resolution ex-vivo MRI template aligned with a group RS-fMRI functional connectivity map of the SC (https://www.marmosetbrainconnectome.org, Schaeffer et al., 2022). This group RS-fMRI map is based on over 70 hours of RS-fMRI collected at ultra-high fields from 31 awake adult marmosets. Marmosets then underwent a second aseptic surgical procedure in which a microdrill (Foredom SR series, Blackstone Industries LLC, Bethel CT) was used to open burr holes of roughly 3mm diameter over the region of PPC identified as described above. This corresponded to approximately to the stereotaxic location of 1.4mm anterior, 6mm lateral indicated for area LIP in the marmoset stereotaxic atlas of Paxinos et al. (2012), and explored in a previous microstimulation study in our lab (Ghahremani et al., 2019). As in that study, we were additionally able to visually identify a small blood vessel and shallow sulcus thought to be homologous to the intraparietal sulcus of macaque. The sites were then sealed with a silicone adhesive (Kwik Sil, World Precision Instruments, Sarasota, FLA, USA) which served to prevent infection and reduce growth of granulation tissue on the dural surface. This seal was removed prior to and replaced following recording sessions after thorough flushing and cleaning of the trephinations.

### Electrophysiological recordings

Recordings were conducted using Neuropixels 1.0 NHP short probes (Jun et al., 2017). The external reference and ground were bridged in all recordings. All recordings were referenced to the reference contact at the tip of the electrode. Data were recorded in two streams, a spike stream sampled at 30 kHz and high-pass filtered at 300 Hz, and an LFP stream sampled at 2.5kHz and low-pass filtered at 300 Hz. Custom Neuropixels electrode holders designed to interface with the dovetail structures on metal cap of the probe base were used with Narishige Stereotaxic Manipulators (SM-25A and SMM-200) to manipulate electrodes for all recordings. IMEC headstages were used with a PXIe-8381 acquisition module, the PXIe-1082 chassis and the MXIe interface were used for data acquisition. 8-bit digital event signals emitted by CORTEX or Monkeylogic and calibrated analog signals for the horizontal and vertical eye positions were recorded using the PXI-6133. Neural and auxiliary signals were synchronized by a TTL pulse emitted by CORTEX or Monkeylogic at target onset. All data was acquired using the SpikeGLX application (v20190413-phase3B2, Karsh, 2019).

For each recording session, we removed the chamber cap and cleaned the recording chamber and dural surface to mitigate the risk of infection. First, we cleaned the outside of the chamber with sterile gauze soaked with 70% isopropyl alcohol solution. The silicone adhesive sealing the trephination was then removed and the dural surface was first flushed with sterile saline delivered via a syringe with a sterile catheter tip. Saline filling the chamber was absorbed with sterile gauze between flushing bouts. In the early stages of the experiments, we tested the use of a suction pump with a sterile tip for this process, but the pump noise caused considerable restlessness in the animals. A 10% iodine solution was then applied, and the area was scrubbed extensively with sterile swabs. We then repeated saline flushing of the area until the solution appeared clear. Any blood or moisture on the dural surface was removed using absorbent surgical eye spears prior to electrode insertion, to avoid fouling of the electrode contacts. Probes were then advanced through the dura using stereotaxic micromanipulators until neural activity no longer appeared on the tip of the electrode where possible. Electrodes were allowed to settle for 30-45 minutes to minimize drift during the recording session. During this time, the animal’s eye position was calibrated as described above. Then, animals performed the visual target selection task as described above until approximately 50 correct trials were obtained in each of the conditions or 45 minutes had passed. Finally, a visual field mapping paradigm was conducted, in which 0.2° dots were briefly flashed (100-200ms SOA, 0-100ms ISI) in a pseudorandomized manner in an evenly spaced 5 x 5 grid spanning +/- 8° abscissa and ordinate. Animals were not required to fixate during this period, and trials where the eyes were closed or moved within +/- 200ms of stimulus onset were removed from analysis offline.

In total, 26 penetrations were conducted across 22 sessions (8 in Marmoset M, 14 in Marmoset N), where 8 penetrations in Marmoset N were conducted with two Neuropixels probes simultaneously. For these penetrations, two probes were adhered back-to-back using dental adhesive (Bisco All-Bond, Bisco Dental Products, Richmond, BC, Canada) and advanced together using a single electrode holder.

### Semi-automated spike sorting

Data collected in the spike stream were additionally high-pass filtered offline at 300 Hz. Putative single unit clusters were then extracted using Kilosort 2 (Pachitariu et al., 2023) Briefly, a common median filter is applied across channels and a “whitening” filter is applied to reduce correlations between channels and maximize local differences among nearby channels. Following these preprocessing steps, templates are constructed based on some initial segment of the data and adapted throughout session with some accommodation for drift over time. Then clusters are separated and merged as necessary.

Following this process, putative single unit clusters were manually curated using the Phy application (Rossant, 2019). Here, clusters were merged or split on the basis of waveforms, cross-correlations and distributions of spike amplitudes. Following merging and splitting clusters as needed, clusters with consistent waveforms, normally distributed amplitudes, a dip in the autocorrelogram at time 0, and consistently observed throughout the recording session were marked as single units, and all others were marked as multi-unit clusters or noise clusters as appropriate. Single unit clusters where the firing rate across the session was at least 0.5 Hz and at most 1% of interspike intervals (ISIs) were within 1 ms (i.e., short ISIs that fall within the refractory period) were retained for all subsequent analyses. For these neurons, short ISI spikes were discarded.

### Identification of task modulated and target discriminating neurons

Neurons were classified as task modulated if activity 40ms from stimulus onset to 25ms after saccade offset significantly differed from baseline activity (200 ms prior to stimulus onset) on contralateral “single-target” trials or “distractor trials. Significance was assessed using paired samples t-tests for each neuron at an alpha level of .05.

Neurons were classified as target discriminating if activity 50-100 ms following stimulus onset significantly differed ipsilateral and contralateral “distractor” trials. Significance was assessed using independent samples t-tests for each neuron at an alpha level of .05.

### Layer assignment based on spectrolaminar LFP analysis

Layer assignment was done as in previous work, using an established spectrolaminar pattern (Mendoza-Halliday et al., 2023). Powerline artifacts were removed at 60 Hz using a butterworth bandstop filter. As these recordings were referenced to the tip of the electrode, as compared to the surface reference used in the recordings of Mendoza-Halliday and colleagues (2023), to recover the pattern they observed, we subtracted the mean activity in channels visually identified as being above the surface from all other channels. Then, the LFP activity aligned to stimulus onset was extracted and the power spectral density (PSD) was computed for each trial using the multi-taper method (Mitra & Pesaran, 1999). This was then averaged across tapers and trials to obtain the mean PSD for a given penetration. The PSD of adjacent channels was then averaged to obtain the mean PSD at each depth (Figure 2d-e). Following visual inspection, power in the 15-22 Hz range was used for the low frequency range and 80-150 Hz was used for the high frequency range. The crossing point in the power of these ranges across depth was marked as the center of layer IV. Upon visual inspection of the density of neurons anchored to this point and the known thickness of layer IV in marmoset PPC, we assigned neurons found from 200 µm below this point to 300 µm above as being in layer IV. Neurons superficial to this range were assigned to layers II/III and those found deeper to layers V/VI.

### Putative cell type classification using peak-trough widths

We clustered neurons as broad and narrow spiking cells on the basis of peak-trough width, which has been suggested to correspond to putative pyramidal cells and interneurons respectively (Ardid et al., 2015; Hussar & Pasternak, 2012; McCormick et al., 1985; Mitchell et al., 2007). For each neuron, the channel at which the spike amplitude had the largest magnitude was selected. The mean waveform at this channel was upsampled to 1 MHz and interpolated using a cubic spline. For cells where the largest amplitude was a peak, the waveform was inverted to ensure that all waveforms exhibited a negative-going pattern. Then, the duration between this trough and the subsequent peak were computed as the peak-trough widths (see Figure 2f). Neurons with a peak-trough width greater than 300 ms were classified as broad spiking (BS) and those with a peak-trough width smaller than 300 ms were classified as narrow spiking (NS).

### Assessing differences in the timing of stimulus-related and discrimination activity across layers and putative cell classes

To assess the contribution of neurons from different cortical layers and putative cell classes to the stimulus-related and discrimination activity across the population, we employed a generalized additive model (GAM). Here, the odds of a spike at a given point in time are estimated using the time from stimulus onset (as a smooth predictor), the layer the neuron is found in (II/III, IV, or V/VI), and putative cell class (NS or BS), with trial and neuron as random effects. For the stimulus-related activity condition (ipsilateral and contralateral was added as a predictor. For the discrimination activity, condition (preferred and nonpreferred) was added as a predictor, where, for each neuron, the stimulus (target or distractor) which elicited the greatest discharge activity was labelled as preferred. Goodness of fit of models as compared to reduced and null models was assessed using the likelihood-ratio chi-squared test. The time where significant stimulus-related activity first emerged was computed by determining where the 99% CI of the difference smooth between ipsilateral and contralateral trials for the “single-target” condition deviated from 0. Similarly, to determine the time at which neurons first significantly discriminated between target and distractor stimuli, we determined where the 99% CI of the difference smooth between preferred and non-preferred trials for the “distractor” condition deviated from 0.

## Acknowledgements

We thank C. Vander Tuin, W. Froese, H. Pettypiece and K. Faubert for expert technical and surgical assistance, and care of the marmosets. This research was supported by the Canadian Institutes of Health Research grant FRN148365 to S.E. and the Canada First Research Excellence Fund to BrainsCAN.

